# Partial Inhibition of Mitochondrial Complex I Reduces Tau Pathology and Improves Energy Homeostasis and Synaptic Function in 3xTg-AD Male and Female Mice

**DOI:** 10.1101/2020.08.19.258236

**Authors:** Andrea Stojakovic, Su-Youne Chang, Jarred Nesbitt, Nicholas P. Pichurin, Mark A. Ostroot, Eugenia Trushina

**Author notes:** Corresponding author: Eugenia Trushina, Ph.D., Department of Neurology, 200 First Street SW, Rochester, MN 55905.

## Abstract

**Background:** Accumulation of hyperphosphorylated Tau (pTau) protein is associated with synaptic dysfunction in Alzheimer’s disease (AD). We previously demonstrated that neuroprotection in familial mouse models of AD could be achieved by targeting mitochondria complex I (MCI) and activating the adaptive stress response. Efficacy of this strategy on pTau-related pathology remained unknown.

**Objective:** To investigate the effect of specific MCI inhibitor tricyclic pyrone compound CP2 on pTau levels, memory function, long term potentiation (LTP), and energy homeostasis in 18-month-old 3xTg-AD mice and explore the potential mechanisms.

**Methods:** CP2 was administered to male and female 3xTg-AD mice from 3.5 - 18 months of age. Cognitive function was assessed using the Morris water maze test. Glucose metabolism was measured in periphery using a glucose tolerance test and in the brain using fluorodeoxyglucose F18 positron-emission tomography (FDG-PET). LTP was evaluated using electrophysiology in the hippocampus. The expression of key proteins associated with neuroprotective mechanisms were assessed by western blotting.

**Results:** Chronic CP2 treatment restored synaptic activity and cognitive function, increased levels of synaptic proteins, improved glucose metabolism and energy homeostasis in male and female 3xTg-AD mice. Significant reduction of human pTau in the brain was associated with increased activity of protein phosphatase of type 2A (PP2A), reduced activity of cyclin-dependent kinase 5 (CDK5) and glycogen synthase kinase 3β (GSK3β).

**Conclusion:** CP2 treatment protected against synaptic dysfunction and memory impairment in symptomatic 3xTg-AD mice, and reduced levels of human pTau, indicating that targeting mitochondria with small molecule specific MCI inhibitors represents a promising strategy for AD.

## Introduction

Alzheimer’s Disease (AD) is an incurable, progressive degenerative disorder characterized by neuronal loss and cognitive dysfunction [1]. Two major hallmarks of AD pathology include the accumulation of amyloid-β (Aβ) peptides in extracellular plaques and a formation of neurofibrillary tangles (NFTs) comprised of hyperphosphorylated tau (pTau) protein [2]. The exact relationship between plaques and NFTs and the severity of neuronal loss and cognitive decline remains unclear. For example, amyloid accumulation in the brain alone does not correlate with the severity of cognitive impairment in patients [3], and the reduction of amyloid plaque load in the brain after immunotherapy did not result in cognitive improvement in AD patients [4]. On the other hand, the increased levels of pTau, the accumulation of the NFTs and the loss of synaptic connectivity correlated the best with cognitive decline and closely reflected the progression of clinical symptoms in AD patients [5-8]. Furthermore, dysfunction in synaptic activity directly associates with a reduced rate of cerebral glucose metabolism detected using FDG-PET, which is prominent in patients with mild cognitive impairment, a prodromal stage of AD [9, 10]. Nevertheless, multiple studies support the notion that the presence of Aβ and pTau synergize to impair the functional integrity of neural circuits contributing to cognitive impairment in AD [11]. Therefore, strategies for neuroprotection should mitigate the pathological phenotypes associated with both Aβ and pTau.

Tau is a microtubule-associated protein that is mainly expressed in neurons. Tau plays an important role in synaptic function regulating microtubule stability and axonal transport [12]. Recent findings present evidence that under physiological conditions, and in response to synaptic activity, tau is recruited to the dendritic spines where it interacts with actin cytoskeleton and actively participates in dendritic spine remodeling [13]. The presence of tau at synapses implicates its involvement in neuronal signaling. Indeed, it was shown that at dendritic spines tau interacts with the Src family non-receptor-associated tyrosine kinase Fyn stabilizing receptor complexes at the post-synaptic density (PSD) [14]. Fyn phosphorylates the GluN2B subunit of N-methyl-D-aspartate receptor (NMDAR) at Tyr1472 facilitating the interaction of the GluN2B complex with the PSD protein 95 (PSD95) [15]. Similarly, tau and PSD95 were shown to contribute to intra-dendritic trafficking and anchoring of amino-3-hydroxy-5-methyl-4-isoxazolepropionic acid receptors (AMPAR) [16, 17]. The GluA1 subunit of AMPAR has several phosphorylation sites including Ser831 and Ser845 that regulate the conductivity of the ion channel and its translocation to the surface membrane [18]. Mice that carry mutations in both Ser831 and Ser845 of GluA1, which prevent phosphorylation of these sites *in vivo*, display defects in spatial learning and memory tasks [19]. Both NMDAR and AMPAR play a central role in the mediating the excitatory synaptic transmission and LTP in the central nervous system (CNS). Thus, the loss of tau function and its aberrant association with the interacting partners could directly affect the integrity of the dendritic spines, synaptic transmission and cognitive function.

In the course of AD progression, tau gets increasingly phosphorylated at several sites (e.g., Thr231, Ser202/Thr205) by multiple kinases, including CDK5 and GSK3β [2, 20]. This hyperphosphorylation results in the accelerated dissociation of pTau from microtubules and the enhanced translocation to the dendritic spines [21]. Abnormal accumulation of pTau to the spines impairs glutamatergic synaptic transmission by reducing the number of functional AMPARs and NMDARs on the surface of the neuronal membranes [22]. In particular, pTau interferes with anchoring of the glutamate receptors to the PSD complex affecting the receptor trafficking to the membrane [22]. The stability of dendritic spines was also compromised by the endocytosis of synaptic AMPAR, leading to spine loss [23, 24]. The disrupted interaction between tau and Fyn further contributes to the destabilization of the dendritic spines and aberrant synaptic function [14]. Along with tau hyperphosphorylation associated with increased activity of CDK5 and GSK3β, deficient pTau dephosphorylation appears to exacerbate the AD pathology. PP2A is one of the major phosphatases that dephosphorylates pTau in mammalian brain [25]. Multiple alterations in PP2A regulation and catalytic activity together with variations in the expression and methylation and/or phosphorylation patterns of its subunits, have been found in AD-affected brain regions [26]. PP2A could directly bind and dephosphorylate pTau at several sites including Ser202/Thr205, Thr231, Ser199, among others [27-29]. PP2A also interacts with protein kinases linked to AD pathogenesis (e.g., GSK3β and CDK5) and neuronal receptors (e.g., NMDAR) [30, 31]. Altered PP2A activity has been directly linked to tau hyperphosphorylation, amyloidogenesis and synaptic deficit in multiple neurodegenerative disorders [26]. In AD, increased inhibitory phosphorylation of PP2A at Tyr307 has been found in tau-rich, tangle-bearing neurons from post-mortem brain tissue [32]. Thus, mechanisms associated with increased pTau accumulation in AD include altered activity of both tau-related kinases and phosphatases.

Currently, there are no disease-modifying strategies for AD. The handful of approved treatments act by counterbalancing altered neurotransmitter activity in symptomatic stage of the disease and produce limited benefits only in a subset of patients. To date, all approaches focused on the amyloid production or clearance failed clinical trials with the attention being shifted to other targets including pTau [33, 34]. However, similar to the failure of Aβ-targeted therapies, strategies to tackle pTau accumulation using immunotherapy or targeting the GSK3β or CDK5 kinase activity did not produce significant clinical outcomes [35-41]. Unexpectedly, one of the approaches shown to improve multiple health parameters in the preclinical models of AD and AD patients includes the induction of the adaptive stress response that allows to acclimate to and overcome stress stimuli [42]. The examples include a non-pharmacological strategies such as exercise [43] or caloric restriction [44, 45] that induce mild energetic stress. While mechanisms of the adaptive stress response are complex, AMP-activated protein kinase (AMPK)-mediated signaling was directly linked to the improvement in cell metabolism, mitochondrial dynamics and function, reduced inflammation and oxidative stress, enhanced autophagy and protein turnover leading to reduced levels of pTau and Aβ [46, 47]. We have shown that similar cascade could be initiated by targeting mitochondrial complex I (MCI) with small molecules that mildly but specifically inhibit MCI [48, 49]. Using multiple transgenic mouse models of familial AD (FAD) that express mutant human amyloid-β protein precursor (AβPP) and/or presenilin 1 (PS1), we demonstrated efficacy of chronic application of the tool MCI inhibitor, a tricyclic pyrone compound (code name CP2), in blocking neurodegeneration, restoration of cognitive and motor phenotype, improved LTP and dendritic spine morphology, reduced levels of Aβ, markers of oxidative stress and inflammation when treatments in independent cohorts of mice were initiated at pre- or symptomatic stages of the disease [48, 49]. Importantly, this strategy improved glucose uptake and utilization in the brain leading to the restoration of energy homeostasis and synaptic function. However, FAD mice utilized in the previous studies did not express human tau. Therefore, it remained unclear whether this innovative therapeutic approach could be efficacious in the presence of accelerated pTau accumulation that synergizes Aβ toxicity. Here we show that CP2 treatment improves cognitive function and synaptic activity in male and female 3xTg-AD mice that along with the mutant human AβPP and PS1, express mutant human tau (P301L) [50]. Chronic treatment resulted in reduced levels of pTau, normalized activity of PP2A, CDK5 and GSK3β, improved LTP, synaptic function, and brain energy homeostasis indicating that targeting mitochondria with small molecules specific MCI inhibitors represent a promising strategy for AD.

## Materials and Methods

All experiments with mice were approved by the Mayo Clinic Institutional Animal Care and Use Committee (IACUC # A0000180-16-R18 and A00001186-16-R18) in accordance with the National Institutes of Health’s Guide for the Care and Use of Laboratory Animals.

### CP2 synthesis

CP2 was synthesized by the Nanosyn, Inc biotech company (http://www.nanosyn.com) as described previously [51] and purified using HPLC. Authentication was done using NMR spectra to ensure the lack of batch-to-batch variation in purity. Standard operating procedures were developed and followed for CP2 stock preparation, storage, cell treatment, and administration to animals.

### Mice

The following female and male mice were used in the study: triple transgenic 3xTg-AD (B6;129-Tg(APPSwe,tauP301L)1Lfa Psen1tm1Mpm/Mmjax) [50] and their appropriate controls B6129SF2/J (F2) (Stock No: 101045, The Jackson laboratory). Genotypes were determined by PCR as described in [52]. All animals were kept on a 12 h–12 h light-dark cycle, with regular feeding and cage-cleaning schedule. Mice were randomly selected to study groups based on their age and genotype. Mice were housed 5 *per* cage, water consumption and weight were monitored weekly. The number of mice in each group was determined based on the 95% of chance to detect changes in 30 - 50% of animals. The following exclusion criteria were established: significant (15%) weight loss, changing in the grooming habits (hair loss), pronounced motor dysfunction (paralyses), or other visible signs of distress (unhealed wounds).

### Chronic CP2 treatment in pre-symptomatic 3xTg-AD mice

The male and female 3xTg-AD and their B6129SF2/J controls (referred to as non-transgenic (NTG) mice) (*n* = 15 *per* group) were given either CP2 (25 mg/kg/day in 0.1% PEG dissolved in drinking water *ad lib*) or vehicle-containing water (0.1% PEG) starting at 3.5 months of age as we described in [48]. Independent groups of mice were continuously treated for 14 months until the age of 18 months. Behavior tests were conducted 3 and 12 months after the beginning of CP2 treatment. *In vivo* FDG-PET and electrophysiology were performed 12 months after the beginning of treatment. After mice were sacrificed, brain tissue was subjected to western blot analysis.

### Morris water maze (MWM) test

Behavioral test was carried out in the light phase of the circadian cycle with at least 24 h between each assessment as we described previously [48]. Spatial learning and memory were investigated by measuring the time it took each mouse to locate a platform in opaque water identified with a visual cue above the platform. The path taken to the platform was recorded with a camera attached above the pool. Each mouse was trained to find the platform during four training sessions *per* day for three consecutive days. For each training session, each mouse was placed in the water facing away from the platform and allowed to swim for up to 60 sec to find the platform. Each training session started with placing a mouse in a different quadrant of the tank. If the mouse found the platform before 60 sec have passed, the mouse was left on the platform for 10 sec and then returned to its cage. If the animal had not found the platform within the 60 sec, the mouse was manually placed on the platform and left there for 10 sec before returning to its cage. A day of rest followed the day of formal testing.

### Intra-peritoneal (IP) glucose tolerance test (IPGTT)

The glucose tolerance test measures glucose clearance after it was delivered using IP injection. Mice were fasted for approximately 16 h, and fasted blood glucose levels were determined before a solution of glucose was administered by the IP injection. Subsequently, blood glucose level was measured at different time points during the following 2 h (30, 60, 90 and 120 min after the IP injection). Mice were injected with 20% glucose solution based on the body weight (2 g of glucose/kg body weight).

### *In vivo* FDG-PET

Mice were fasted one hour prior to the IP injection of 270 uCi of FDG in 200 µL injection volume prepared the same day at the Mayo Clinic Nuclear Medicine Animal Imaging Resource. Imaging was conducted 30 min post injection. Prior to imaging, mice were individually placed in an anesthesia machine (Summit Medical Equipment Company, Bend, Oregon) and anesthetized with 4% isoflurane with 1-2 LPM oxygen. Anesthesia was further maintained with 2% isoflurane delivered by a nose cone. Mice were placed in MicroPET/CT scanner (Scanner Inveon Multiple Modality PET/CT scanner, Siemens Medical Solutions USA, Inc.). Mouse shoulders were positioned in the center of the field of view (FOV), and PET acquisition was performed for 10 min. CT scanning parameters were as following: 360 degree rotation; 180 projections; Medium magnification; Bin 4; Effective pixel size 68.57; Tranaxial FOV 68 mm; Axial FOV 68 mm; 1 bed position; Voltage 80 keV; Current 500 uA; Exposure 210 ms. CT reconstruction parameters: Alogorithm: Feldkamp, Downsample 2, Slight noise reduction with application of Shepp-Logan filter. Final analysis was done using PMOD Biomedical Image Quantification and Kinetic Modeling Software, (PMOD Technologies, Switzerland). The volume of interest (VOI) was created on the CT image of the entire brain. The VOI was applied to the corresponding registered PET scan. The volume statistics were recorded. The percentage of brain glucose uptake was calculated by correcting the measured concentration of uCi recorded after 30 min to the amount of injected dose.

### Electrophysiology

Female vehicle- and CP2-treated 3xTg-AD mice 18 months of age treated for 14 months (n = 4 *per* group), were used for electrophysiological analysis. Mice were deeply anesthetized with isoflurane and perfused with cold artificial cerebrospinal fluid (ACSF), where NaCl was substituted with sucrose to avoid excitotoxicity. Brains were quickly removed and transferred into a cold slicing solution containing an ACSF. Transverse slices were made at 300 - 350 µm thickness using a vibratome (VT-100S, Leica). Slices were incubated in ACSF containing 128 mM NaCl, 2.5 mM KCl, 1.25 mM NaH2PO4, 26 mM NaHCO3, 10 mM glucose, 2 mM CaCl2, and 1 mM MgSO4, aerated with 95% O2/5% CO2. Slices were maintained at 32 °C for 13 min, and then maintained at room temperature throughout the entire experiment. For electrophysiological recording, each slice (2 - 3 slices *per* mouse) was transferred to a recording chamber, and ACSF was continuously perfused at the flow rate of 2 - 3 ml/min. A single recording electrode and a single bipolar stimulation electrode were placed on top of the slice. A boron-doped glass capillary (PG10150, World Precision Instruments) was pulled with a horizontal puller (P-1000, Sutter Instrument) and filled with ACSF for extracellular recording. Under the microscope (FN-1, Nikon), the recording electrode was placed in the CA1 area of the hippocampus. The bipolar stimulation electrode (FHC) was placed at the Schaffer collaterals. The distance between two electrodes was over ∼200 µm. To define a half response of stimulation, various intensities of electrical stimulation were applied (10 - 500 µA). However, the pulse width was fixed at 60 µsec. Once the stimulation parameter was determined to generate a half maximum of evoked field excitatory postsynaptic potential (fEPSP), this stimulation intensity was used for the LTP experiments. To measure LTP, test stimulation was applied every 30 sec for 30 min to achieve a stable baseline. Once the stable baseline was achieved, a tetanic stimulation (100 Hz for 1 sec) was applied three times at 30 sec intervals. Initial slopes of the fEPSP were used to compare synaptic strength. The slopes of the fEPSP were analyzed by pCLAMP10.5.

### Western blot analysis

Levels of proteins were determined in the cortico-hippocampal region of the brain of vehicle- and CP2-treated male and female 3xTg-AD mice (*n* = 4 mice *per* group) using western blot analysis. Tissue was homogenized and lysed using 1× RIPA buffer plus inhibitors. Total protein lysates (25 µg) were separated in equal volume on 4 – 20% Mini-PROTEAN TGX™ Precast Protein Gels (Bio-Rad, cat. # 4561096) and transferred to an Immun-Blot polyvinylidene difluoride membrane (PVDF cat. # 1620177). The following primary antibodies were used: phospho-AMPK (Thr 172) (1:1000, Cell Signaling Technology, cat. # 2535), AMPK (1:1000, Cell Signaling Technology, cat. #2532), Synaptophysin (1:200, Santa Cruz Biotechnology, Santa Cruz, CA, cat. # 17750), PSD95 (1:1000, Cell Signaling Technology, cat. # 2507), GluA1 (1:1000, Cell Signaling Technology, cat. # 13185), phospho-AMPAR 1 (Ser845) (1:1000, Cell Signaling Technology, cat. # 8084), GluN2B (1:1000, Cell Signaling Technology, cat. # 14544), phospho-NMDAR 2B (Tyr1472) (1:1000, Cell Signaling Technology, cat. # 4208), AT8 (1:1000, Thermo Fisher, cat. # MN1020), AT180 (1:1000, Thermo Fisher, cat. # MN1040), HT7 (1:1000, Thermo Fisher, cat. # MN1000), Fyn (1:1000, Cell Signaling Technology, cat. # 4023), phosphor-Src (Tyr416) (1:1000, Cell Signaling Technology, cat. # 6943), CDK5 (1:1000, Cell Signaling Technology, cat. #2506), p35/p25 (1:1000, Cell Signaling Technology, cat. # 2680), GSK3β (1:1000, Cell Signaling Technology, cat. #9832), phospho-GSK3β (Ser9) (1:1000, Cell Signaling Technology, cat. #9322), PP2Ac (1:1000, Cell Signaling Technology, cat. #2038), phospho-PP2Ac (Tyr 307) (1:1000, Santa Cruz, cat. #sc-12615), Tubulin (1:5000, Biovision, cat. #3708), β-Actin (1:5000, Sigma-Aldrich, cat. # A5316). The following secondary antibodies were used: donkey anti-rabbit IgG conjugated with Horseradish Peroxidase (1:10000 dilution, GE Healthcare UK Limited, UK) and sheep anti-mouse IgG conjugated with Horseradish Peroxidase (1:10000 dilution, GE Healthcare UK Limited, UK). Band quantification was done using Image LabTM v. 6.0.

### Statistics

The statistical analyses were performed using the GraphPad Prism (Version 8, GraphPad Software, Inc., La Jolla, Ca). Statistical comparisons among four groups in the MWM test were analyzed by one or two-way ANOVA. The Fisher’s LSD *post hoc* analysis was used if significant interaction among groups was found. Significant differences between vehicle and CP2-treated groups of the same sex were analyzed by Student *t*-test. Data are presented as mean ± SEM for each group of mice.

## Results

### Long-term CP2 treatment ameliorates cognitive decline in male and female 3xTg-AD mice

The tricyclic pyrone compound CP2 penetrates the blood-brain barrier, has good oral bioavailability, and does not induce any toxicity at doses up to 50 mg/kg/day [48, 49]. CP2 accumulates in mitochondria where it competes with the flavin mononucleotide (FMN) for binding to the MCI mildly inhibiting its activity [48, 53]. To ascertain if chronic treatment averts the progression of cognitive impairment, pre-symptomatic male and female 3xTg-AD mice were treated with CP2 (25 mg/kg/day in drinking water *ad lib*) from 3.5 until 18 months of age (Fig.1A). Non-transgenic F2 (NTG) male and female age-matched mice were used as controls. Consistent with our previous studies [48, 49], CP2 treatment in 3xTg-AD mice was safe and did not cause noticeable side effects. Cognitive function was evaluated 12 months after the beginning of treatment using the MWM test, a well-validated paradigm for testing spatial learning abilities commonly utilized in characterization of memory dysfunction in 3xTg-AD mice [54]. Consistent with the previous reports, 16-month-old vehicle-treated male and female 3xTg-AD mice had impaired learning acquisition compared to NTG mice (Fig.1B,E). CP2 treatment improved spatial learning and memory in male and female 3xTg-AD mice based on the performance over four days of training demonstrating significantly reduced time to find the platform compared to untreated counterparts (Fig.1B,E). Cognitive improvement persisted over time where CP2-treated 3xTg-AD mice of both sexes showed reduced latency to find the location of a hidden platform 24 h after the last training session (Fig.1C,F). The analysis of the animal meandering, which represents the number of changes in the swimming direction done by a mouse during the process of searching for the hidden platform, showed that CP2-treated 3xTg-AD mice were more focused in their attempt to find the platform compared to vehicle-treated mice (Fig.1D,G). CP2 treatment was most efficacious in 3xTg-AD mice and did not significantly affect the performance of NTG mice. These observations suggest that long-term chronic treatment with MCI inhibitor CP2 improves cognitive function in male and female 3xTg-AD mice.

### Chronic CP2 treatment improves glucose homeostasis in the brain and periphery in 3xTg-AD mice

The 3xTg-AD mice develop brain hypometabolism measured using FDG-PET and peripheral glucose intolerance at 16 months of age recapitulating prominent mechanisms of human disease [55, 56]. To determine whether CP2 treatment improves cerebral and peripheral glucose metabolism, we first measured glucose uptake in the brain of 3xTg-AD mice *in vivo* using translational biomarker FDG-PET, a well-established method that provides differential diagnosis of AD patients [57]. Results confirmed that male and female vehicle-treated 3xTg-AD mice developed significant cerebral glucose hypometabolism at 16 months of age (Fig.2A-D). CP2 treatment for 12 months restored glucose uptake in the brain of both male and female 3xTg-AD mice to the levels observed in NTG age- and sex-matched control mice (Fig.2A-D). We next examined the effect of treatment on peripheral glucose metabolism by applying the IPGTT to 3xTg-AD mice treated with CP2 for 12 months. We found that CP2-treated female 3xTg-AD mice had significantly improved glucose tolerance compared to vehicle-treated littermates (Fig. 2E). CP2-treated male 3xTg-AD mice showed a trend toward an improvement in comparison to their vehicle-treated counterparts (Fig.2F). These data demonstrate the overall improvement in glucose metabolism in 3xTg-AD mice treated with MCI inhibitor CP2.

**Fig. 1.**
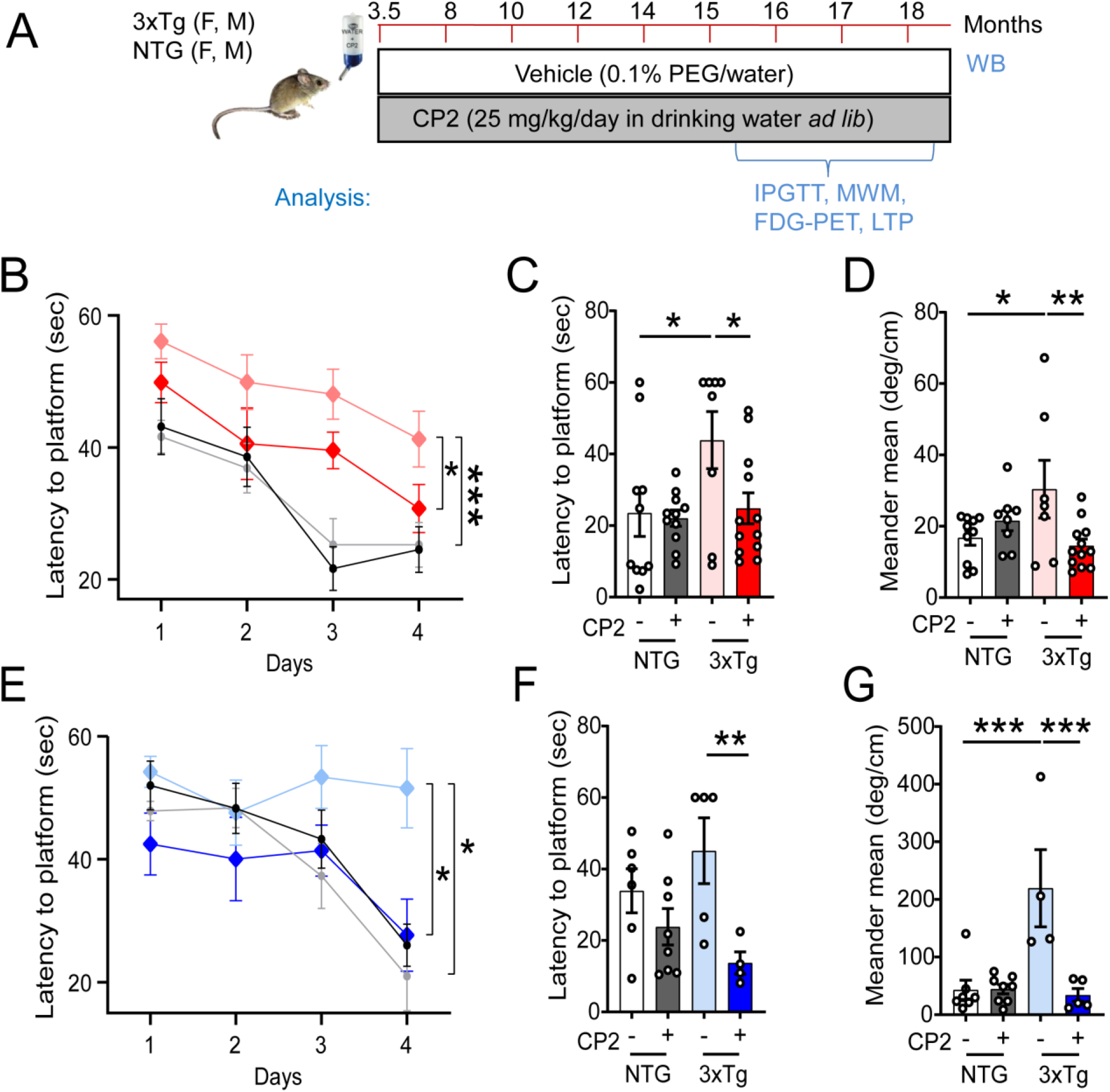
Long-term CP2 treatment ameliorates cognitive deficit in male and female 3xTg-AD mice. **A)** The timeline of CP2 treatment in 3xTg-AD and NTG male and female mice (*n* = 15 *per* each group). **B)** Memory acquisition was done based on the time of escape latency established over 4 days of training in the MWM test. **C)** Latency to reach the platform 24 h after the last training session showed that CP2 treatment significantly improved memory retention in female 3xTg-AD mice. **D)** CP2 treatment significantly reduced meandering (the direction changes) in female 3xTg-AD mice. **E)** CP2 improved memory acquisition in 3xTg-AD male mice, whereas vehicle-treated 3xTg-AD mice had impaired learning acquisition compared to NTG mice. **F)** Latency to reach the platform 24 h after the last training session showed that CP2 treatment significantly improved memory retention in male 3xTg-AD mice. **G)** CP2 treatment significantly reduced meandering (the direction changes) in male 3xTg-AD mice. Differences between individual groups were analyzed by two-way ANOVA with Fisher’s *post-hoc* test. Data are presented as mean ± SEM. *P<0.05, **P<0.01, *** P<0.001. Light blue and light red – vehicle-treated 3xTg-AD male and female mice, respectfully; dark blue and dark red - 3xTg-AD+CP2 male and female mice, respectfully; grey - NTG+CP2 (female mice in C and D; male mice in F and G); black – vehicle-treated NTG (female mice in C and D; male mice in F and G); *n* = 10-15 mice *per* group.

**Fig 2.**
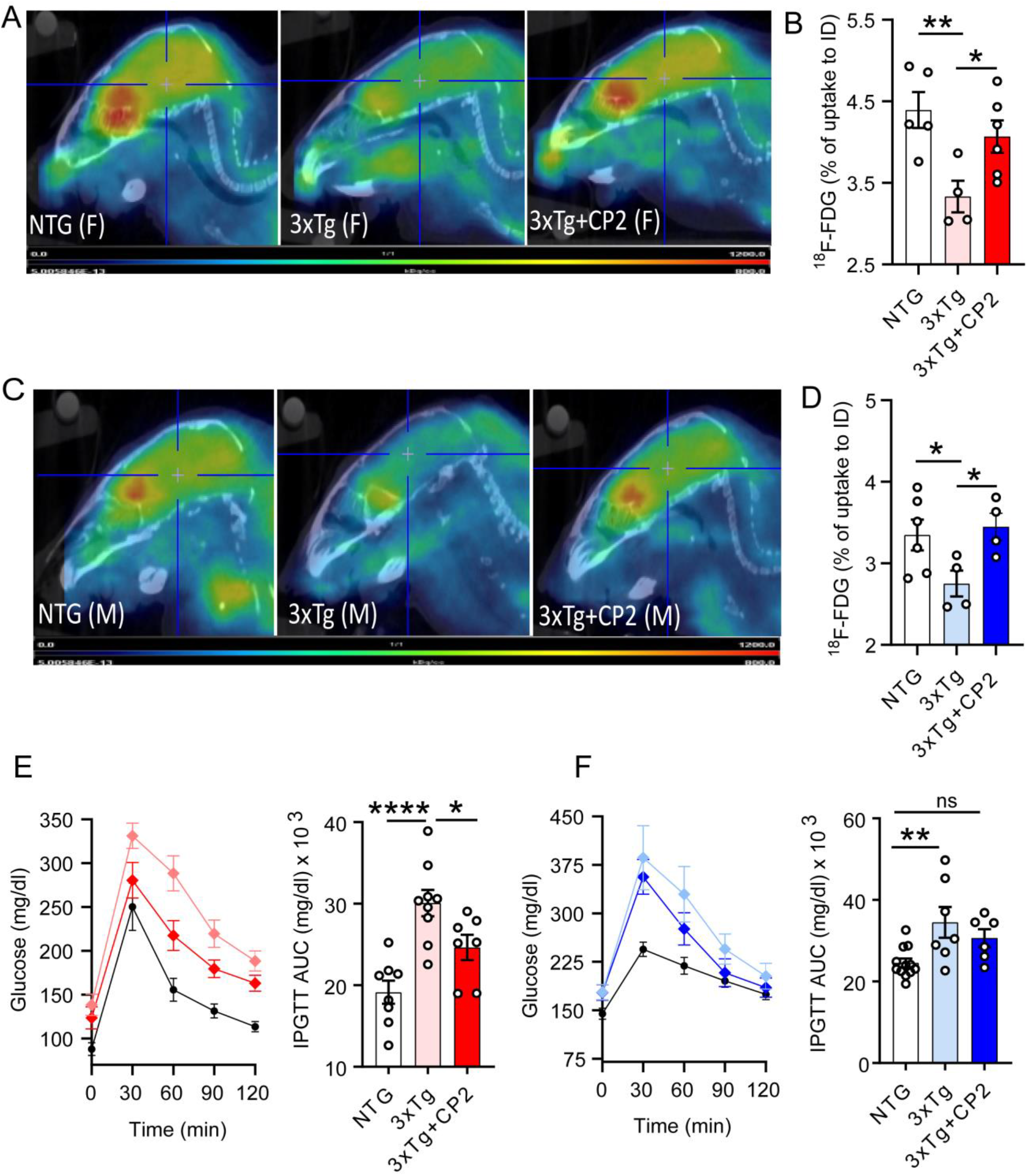
Chronic CP2 treatment improves glucose homeostasis in the brain and periphery in 3xTg-AD mice. **A**,**C)** Representative images of the ^18^F-fluorodeoxyglucose (^18^F-FDG) uptake in the brain of CP2-treated female (**A**) and male (**C**) 3xTg-AD mice compared to vehicle-treated age- and sex-matched 3xTg-AD and NTG mice measured using *in vivo* FDG-PET 14 months after the beginning of treatment. **B**,**D)** Quantification of glucose uptake measured using *in vivo* FDG-PET from (**A** and **C**) as a percent of changes compared to the initial dose (ID). *N* = 5 mice *per* each group. **E)** CP2 treatment increases glucose tolerance in 3xTg-AD female mice measured by IPGTT test. *N* = 6-12 mice *per* group. **F)** CP2 treatment did not significantly improved glucose tolerance in 3xTg-AD male mice. *N* = 6-12 mice *per* group. Differences between individual groups were analyzed by one-way ANOVA with Fisher’s *post-hoc* test. Data are presented as mean ± SEM. *P<0.05, **P<0.01, **** P<0.0001.

### CP2 treatment improves LTP and synaptic function in 3xTg-AD mice

Cerebral glucose hypometabolism and cognitive dysfunction are related to structural and functional impairment of dendritic spines [58]. Therefore, we next explored whether the improvement in memory and brain energy homeostasis observed in CP2-treated 3xTg-AD mice were related to the augmented function of pyramidal neurons in the CA1 region of the hippocampus (Fig.3A). LTP in the CA1 region represents a form of plasticity responsible for learning and memory, and it critically relies on the activation of NMDARs for its induction and increased insertion of AMPARs for its expression [59]. Previous studies demonstrated that significant reduction of the LTP was already prominent in 12-months-old 3xTg-AD mice [56]. To determine fEPSP, tetanic stimulation was applied to Schaffer collaterals-CA1 to induce and record LTP over 60 min. In these experiments, we utilized only female mice. Significant current associated with strong LTP was recorded in CP2-treated 3xTg-AD mice for 30 min, whereas vehicle-treated 3xTg-AD mice did not exhibit significant LTP formation (Fig.3B,C). A three-fold increase in the LTP formation 30 min after the stimulation was achieved in CP2-treated 3xTg-AD mice, whereas the response in vehicle-treated mice was almost nonexistent (Fig.3B,C).

Previous findings in 3xTg-AD mice revealed a significant decrease in the phosphorylation of multiple AMPAR and NMDAR subunits [60]. In comparison to those results, we found that CP2 treatment significantly increased phosphorylation of GluA1 subunit of AMPAR at Ser845 and phosphorylation of NMDAR subunit GluN2B at Tyr1472 (Fig.3D,E). These data demonstrate that functional alterations in AMPAR and NMDAR associated with synaptic deficit in 3xTg-AD mice were restored by CP2 treatment resulting in increased LTP. Improved LTP in CP2-treated 3xTg-AD mice was also associated with increased levels of synaptophysin and PSD95 (Fig.3D,E), which could account for improved synaptic transmission in the hippocampus.

### CP2 treatment reduces levels of human pTau in 3xTg-AD mice

We next determined whether CP2-mediated improvement of LTP, synaptic function and a performance in the MWM were associated with the reduction of pTau levels using western blot analysis. In both male and female 3xTg-AD mice, CP2 treatment significantly reduced pTau detected using specific antibodies AT8 (Ser202/Thr205) and AT180 (Thr231) (Fig.4A-D). Levels of total tau detected using HT7 antibody were not affected by the treatment (Fig. 4B,D). Compared to male mice, female CP2-treated 3xTg-AD mice had reduced Fyn levels in the brain, consistent with an improvement in dendritic spine function associated with the restoration of tau-Fyn physiological interactions (Fig.4A-D). We also detected a reduced activity of two main kinases, CDK5 (based on the reduction in levels of p35 and p25 proteins) and GSK3β (increased inhibitory phosphorylation at Ser9), which are directly involved in tau phosphorylation (Fig.4A-D). In male mice, these changes did not reach statistical significance and, apart from GSK3β, were showing trends.

**Fig 3.**
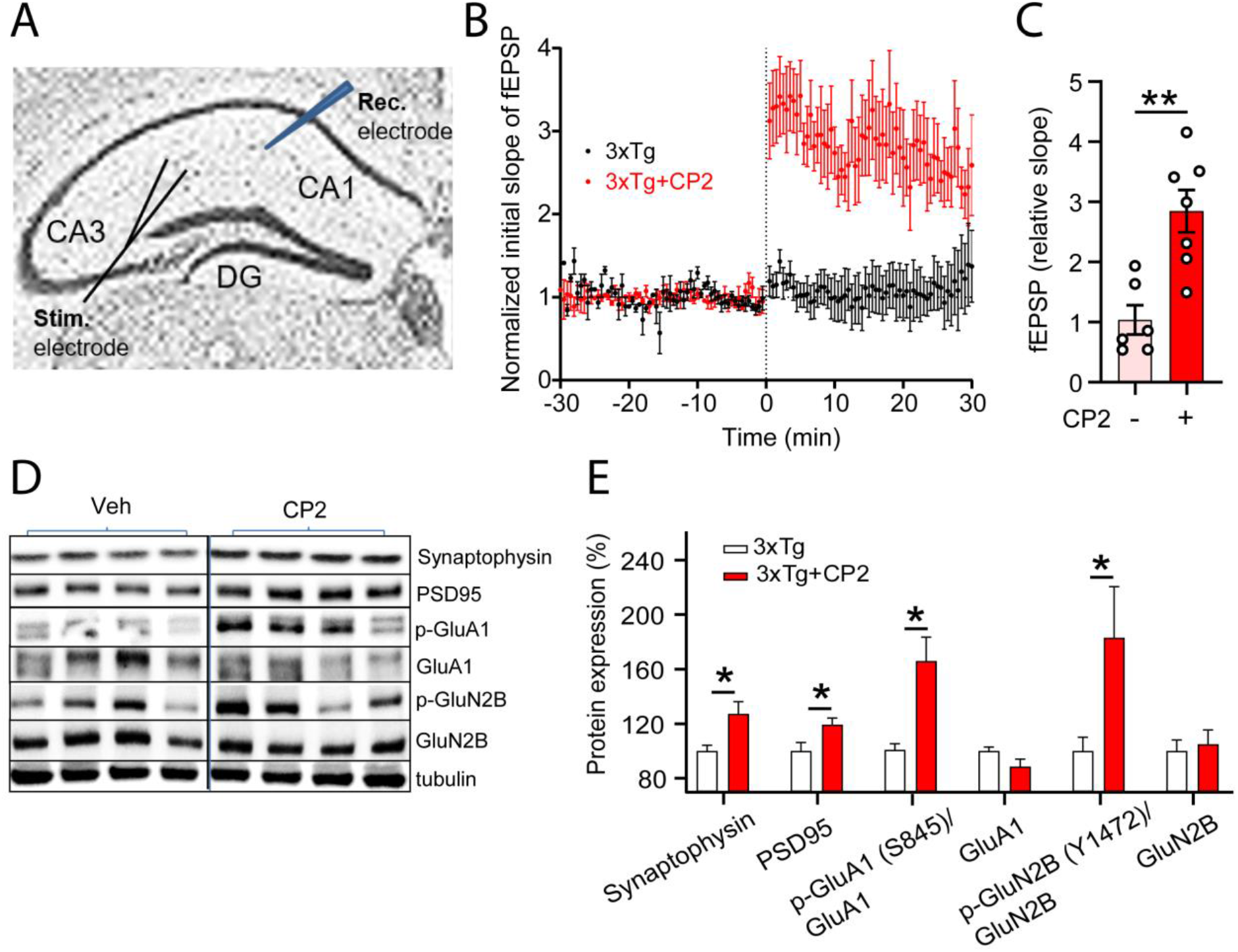
CP2 treatment improves synaptic function and LTP in 3xTg-AD mice. **A)** Experimental setup for stimulation-evoked local field potential (LFP) measurement in the hippocampal acute slice. The stimulation (Stim.) electrode was placed at the Schaffer collaterals and the recording (Rec.) electrode was placed at the striatum radiatum in the CA1 region. To induce LTP, three tetanic stimulations (100 Hz, 60 µsec-pulse width for 1 sec) were applied with 3 sec intervals. In the slice, LTP was induced and maintained over 30 - 50 min. Slope of fEPSPs was measured and results normalized to the average value established during the 30 min baseline period. Recording continued for at least 60 min following a tetanic stimulation and the first 30 min was used to calculate the LTP. **B)** CP2 treatment improves LTP formation in 3xTg-AD female mice. Traces represent mean ± SEM *per* time. *N* = 2-3 slices from 3–5 mice *per* group.**C)** LTP intensity in CP2-treated female 3xTg-AD mice compared to vehicle-treated counterparts at 30 min after the stimulation. *N* = 2 - 3 slices from 3–5 mice *per* group. **D)** Representative western blot analysis of protein markers involved in synaptic function of dendritic spines in the brain tissue of CP2- and vehicle-treated female 3xTg-AD mice. *N* = 4 mice *per* group. **E)** Quantification of western blot data from **(D)** showing a percent of changes in protein expression after CP2 treatment. Significant increase was observed in levels of synaptophysin, PSD95, and GluA1 and GluN2B phosphorylation. No changes were detected in total levels of GluA1 and GluN2B. Differences between individual groups were analyzed by Student *t*-test. Data are presented as mean ± SEM. *P<0.05, **P<0.01.

**Fig 4.**
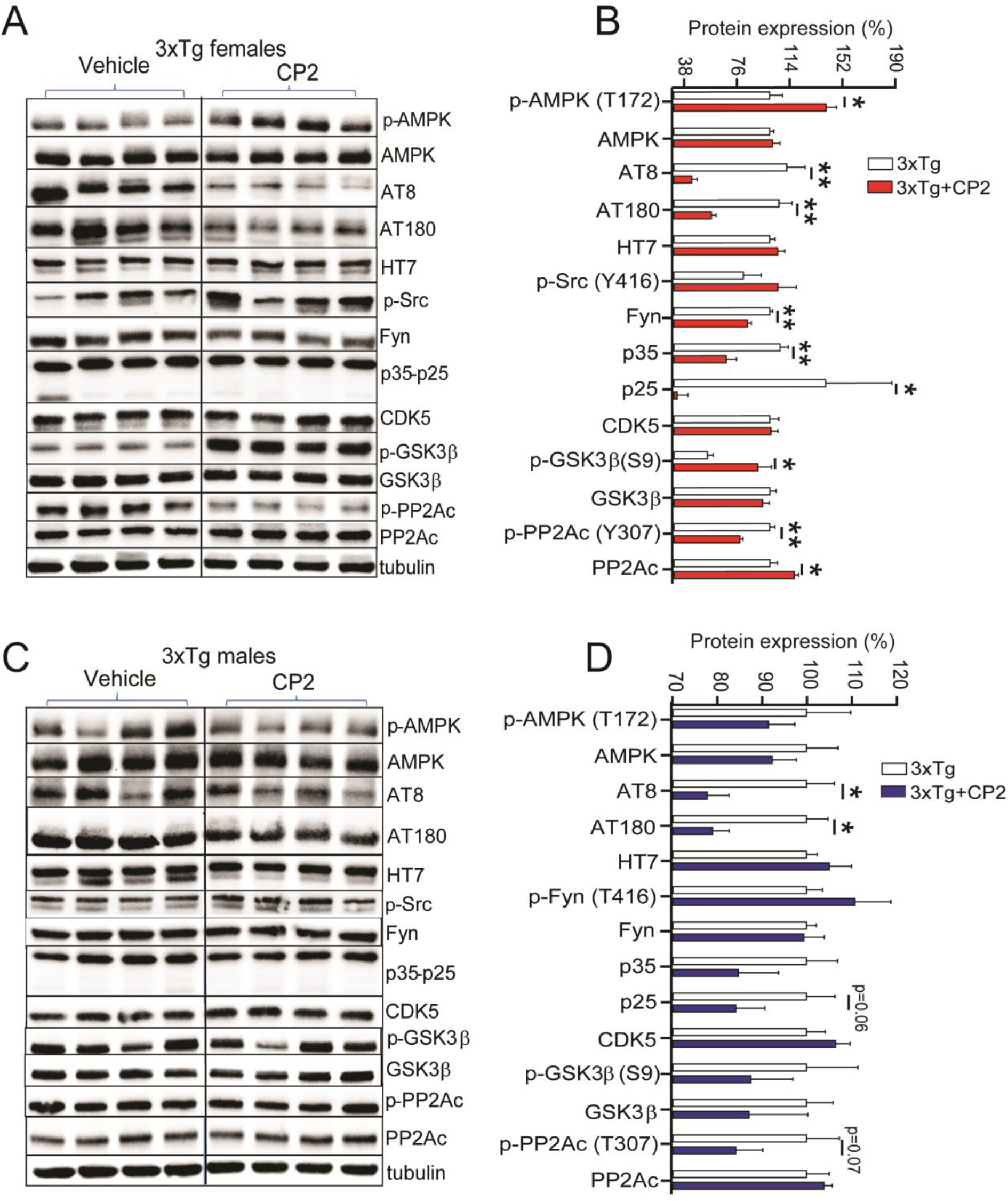
CP2 treatment reduces levels of human pTau in the brain tissue of male and female 3xTg-AD mice. **A**,**C)** Western blot analysis of protein expression in the brain tissue of vehicle- and CP2-treated female (**A**) and male (**C**) 3xTg-AD mice treated for 14 months. **B**,**D)** Changes in protein expression in CP2-treated 3xTg-AD female (**B**) and male (**D**) mice (from **A** and **C**, respectively) relative to the levels in vehicle-treated counterparts. *N* = 4 mice *per* group. Differences between individual groups were analyzed by Student *t*-test. Data are presented as mean ± SEM. *P<0.05, **P<0.01.

Activation of AMPK was implicated in the induction of the adaptive stress response associated with CP2 efficacy [49]. AMPK is also a direct activator of PP2A, the main protein phosphatase involved in pTau de-phosphorylation in mammalian cells [61-63]. We found that levels of pAMPK were significantly increased only in CP2-treated female 3xTg-AD mice (Fig. 4B,D). Consistently, the inhibitory phosphorylation at the Tyr307 of PP2Ac was significantly reduced only in 3xTg-AD female mice while male mice showed a trend (Fig. 4B,D). The total levels of PP2Ac were significantly elevated in CP2-treated female 3xTg-AD mice with males showing a trend. Taken together, these data demonstrate that, at least in females, CP2 treatment activates the AMPK-dependent integrated signaling cascade leading to a reduction of human pTau.

## Discussion

Our previous studies demonstrated that chronic treatment with partial MCI inhibitor CP2 blocked neurodegeneration and improved cognitive and metabolic functions associated with AD in multiple mouse models of FAD that recapitulated primarily the Aβ pathology [48, 49]. The significance of the current findings is the demonstration that CP2 treatment restores pTau-related pathology including abnormal LTP and dendritic spine function in symptomatic 3xTg-AD mice. This mouse model is one of the most relevant animal models of AD, where the disease phenotype is associated with the synergistic toxicity of both Aβ and pTau. The demonstration that CP2 treatment significantly reduced levels of human pTau provides further support for a high translational potential of this innovative mitochondria-targeted therapeutics. The outcomes of this study are in agreement with our previous observations showing that a prolonged chronic administration of MCI inhibitors is safe and efficacious when administered to pre- or symptomatic aged mice without causing side effects. We chronically treated 3xTg-AD mice with CP2 for ∼14 months, where not a single animal was removed from the study due to adverse treatment-related complications. Consistent with the previous findings in APP/PS1, APP and PS1 mice [48, 49], results of this study demonstrate that CP2 treatment attenuated pathological mechanisms, including brain hypometabolism, glucose uptake, and synaptic and cognitive dysfunction, which are prominent in AD patients and are recapitulated in 3xTg-AD mice. Importantly, treatment efficacy in 3xTg-AD mice was shown *in vivo* using translational biomarker FDG-PET currently used as one of the tools to diagnose AD patients.

Additional strength of the current study is in the complementary experimental approach where outcomes of behavior test were supported with *in vitro* and *ex vivo* functional and mechanistic studies providing compelling evidence in support of the conclusions.

The major findings of this study include the demonstration of LTP restoration, cognitive improvement and a reduction of pTau levels in 18-month-old 3xTg-AD mice after chronic treatment with MCI inhibitor. Significant dysfunction in spatial memory and learning is well established in 3xTg-AD mice [54]. CP2 treatment improved a performance of 3xTg-AD mice in the MWM test, a standard test for cognitive assessment. Along with the significantly shortened time to find the platform, CP2-treated mice also had improved attention while performing this task. Spatial memory and learning was improved in male and female 3xTg-AD mice to the levels observed in age- and sex-matched NTG control mice. Synaptic transmission and LTP are essential components of memory formation. The excitatory synapses are formed on dendritic spines, the specialized domains that encompass multiple synaptic proteins, receptors, mitochondria and actin filaments. LTP is triggered by the intense activation of the NMDA receptors producing a signaling cascade that causes the recruitment of AMPA receptors into the postsynaptic membrane. Synaptic strength critically depends on the spine morphology, PSD area, presynaptic active zone area, and the number of AMPARs [64]. Based on the mounting evidence from epidemiological studies, synaptic loss represents the earliest pathological mechanism of AD and is the best predictor of the clinical symptoms of patients with AD [6]. Recent data reveled that along with its function as a microtubule-stabilizing protein, tau also plays a role at synapses. Healthy neurons maintain a spatial gradient of tau protein with greater concentration in axons compared to dendritic compartments [12]. Under physiological conditions, tau participates in synaptic function in dendritic spines by stabilizing NMDAR-PSD95 complex in coordinated interaction with Fyn. In AD, this gradient becomes inverted [65], enabling the accumulation of pTau in dendritic spines leading to synaptic loss [66]. The expression of mutant tau (e.g., P301L) and its increased synaptic localization were associated with concomitant accumulation of Fyn kinase in dendritic spines [14]. Fyn overexpression and localization to the dendritic spines were linked to accelerated cognitive impairment [67], whereas down-regulation of Fyn restored memory function and synaptic density in AD mouse model [67]. Therefore, a reduction of pTau levels should improve the dendritic spine morphology and function and augment LTP. Indeed, we have found that CP2 treatment not only reduced levels of human pTau (AT8, AT180) in male and female 3xTg-AD mice but also levels of Fyn, specifically in female 3xTgAD mice. Lower levels of pTau in neurons should led to the normalized activity and localization of Fyn, and improved dendritic spine function, which was further supported with the demonstration of the LTP restoration in the hippocampus of 3xTg-AD mice.

Further support for CP2-dependent improvement of synaptic function comes from experiments that evaluated changes in phosphorylation of NMDAR and AMPAR subunits. The AMPARs play a central role in the modulation of excitatory synaptic transmission in the CNS, where their function is regulated by the composition and the phosphorylation state of their four subunits [68]. Changes in AMPAR signaling and the loss of dendritic spine morphology and function were identified as a leading cause of early synaptic dysfunction in 3xTg-AD mice [60]. In particular, these mice exhibited the abnormal membrane trafficking and the insertion of hippocampal GluA1-containing AMPARs during LTP stimulation [60]. The impairment in AMPAR activity is largely mediated by a reduction in phosphorylation of its multiple subunits, including Ser845, which affects AMPAR membrane trafficking [60]. In agreement with the findings demonstrating the improved LTP and cognitive function in CP2-treated 3xTg-AD mice, we showed that treatment increased phosphorylation of AMPAR subunits at Ser845 to the levels observed in NTG mice [60], supporting functional increase in synaptic activity in 3xTg-AD mice. Phosphorylation of AMPARs at Ser845 also prevents their sorting to lysosomes for degradation [69], facilitates an increase in mean open time of the receptor [19], and promotes GluA1 delivery to the dendritic spines in order to make AMPARs available for subsequent synaptic recruitment during LTP formation [70]. CP2 treatment also increased the phosphorylation of NMADR at Tyr1472 in 3xTg-AD mice bringing it to the levels observed in NTG mice [60]. This phosphorylation is essential for a proper localization of these receptors to dendritic spines and their function [71, 72]. Since aberrant increase in phosphorylation could contribute to excitotoxicity through increased stabilization of receptor complexes with PSD and over-activation [14], it is important to note that CP2 treatment restored phosphorylation of AMPAR and NMDAR subunits to the levels observed in age-matched NTG controls [60]. Improved LTP in 3xTg-AD mouse brain tissue was associated with increased levels of synaptic proteins PSD95 and synaptophysin, further supporting improvement in synaptic activity.

The major kinases that phosphorylate tau in AD include CDK5 and GSK3β. Our data indicate that beneficial molecular mechanisms engaged by the treatment are upstream of tau kinases and act to reduce their activity, while having no impact on total levels of tau expression. CP2 was profiled against 250 kinases in the selectivity screen, where it demonstrated the lack of kinase activity at 10 μM, a concentration that is ∼ 100 x higher compared to concentrations found in the brain tissue from treated mice [49]. Our previous studies demonstrated that mild MCI inhibition with CP2 induced AMPK activation in neurons *in vitro* and in the brain tissue of FAD mice *in vivo* leading to an induction of an integrated signaling cascade comprised of the mechanisms independently shown as neuroprotective [48, 49]. AMPK-mediated signaling has been directly linked to the regulation of cell metabolism, mitochondrial dynamics and function, inflammation, oxidative stress, protein turnover, Tau phosphorylation, and amyloidogenesis [46]. Combined analysis performed using multiple types of genome-wide data identified a predominant role for metabolism-associated biological processes in the course of AD, including autophagy and insulin and fatty acid metabolism, with a focus on AMPK as a key modulator and therapeutic target [73]. While the molecular mechanisms associated with AMPK activation are complex and require further investigation, AMPK could reduce the activity of GSK3β via activation of Wnt signaling pathway [74]. Interestingly, Wnt activation was also shown to improve brain glucose uptake and utilization providing neuroprotection in a mouse model of AD [75]. Furthermore, we found that CP2 treatment increases the activity of PP2Ac, the most important phosphatase that is capable of dephosphorylating pTau at AD specific phosphosites in the brain [25]. AMPK is a direct activator of PP2A in mammalian cells [36-38]. A reduction of the expression and the activity of PP2A has been found in the brain tissue of AD patients [76] making PP2A a potential target for drug development for AD and related tauopathies. Indeed, we found that CP2 treatment significantly activated PP2Ac in female 3xTg-AD mice concomitant with AMPK activation while in male mice PP2Ac activation did not reach statistical significance producing only a trend. Furthermore, consistent with the role of PP2Ac in regulating the activity of CDK5 and tau phosphorylation in AD, we found a significant reduction of CDK5 activity in CP2-treated female 3xTg-AD mice, while in male mice there was a trend [77]. Taken together, these data demonstrate that partial inhibition of MCI activates beneficial molecular mechanisms that specifically reverse pTau-mediated toxicity in AD restoring LTP and cognitive function in 3xTg-AD mice (Fig. 5).

**Fig 5.**
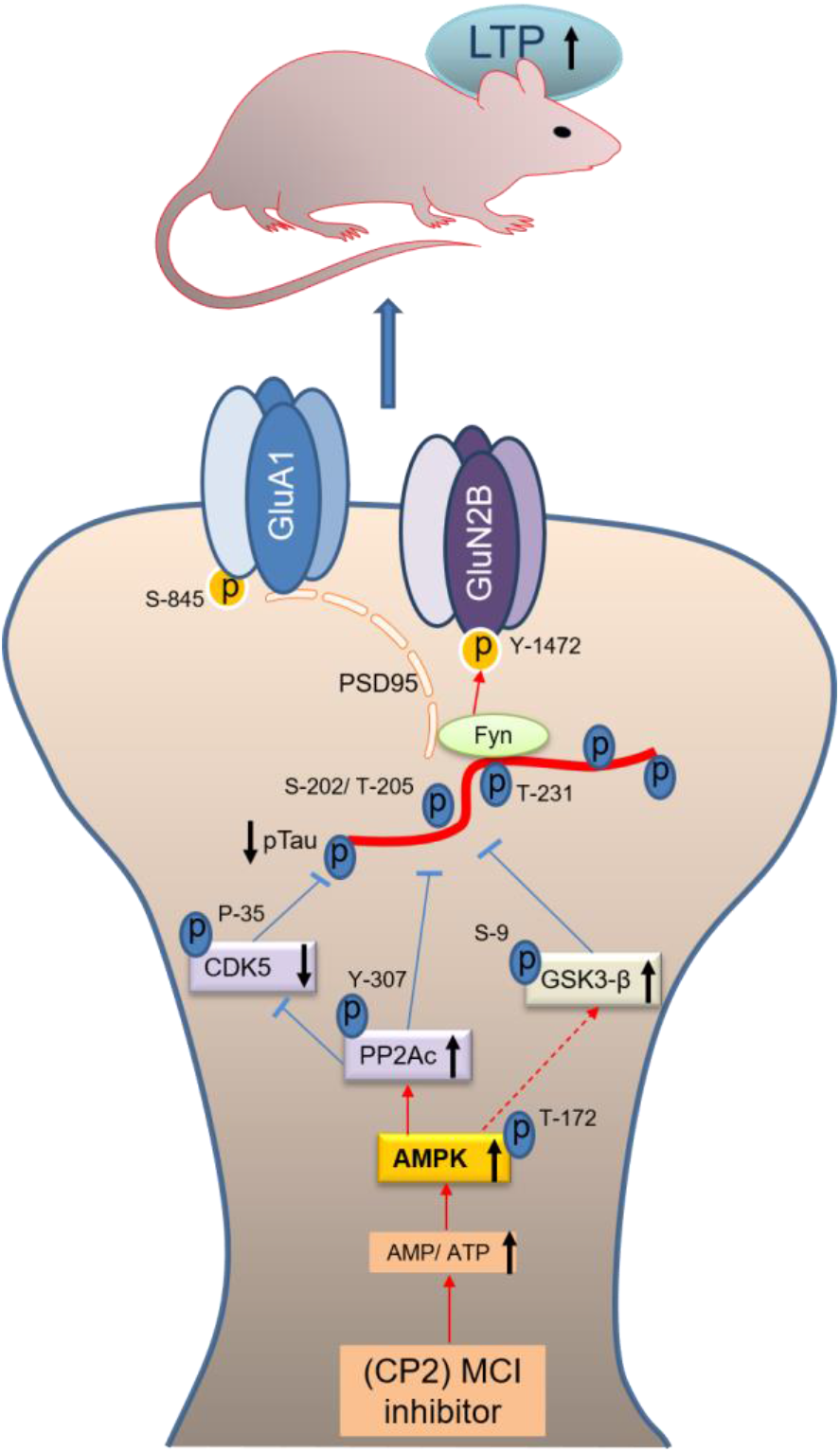
Schematic diagram of the mechanisms involved in CP2-mediated neuroprotection. Mechanisms of CP2-dependent restoration of pTau-mediated synaptic dysfunction. Under physiological conditions, tau transports Fyn to the dendritic spines where Fyn phosphorylates the NR2B subunit of NMDARs at Y-1472 leading to the stabilization of the NMDAR:PSD95 complex. Increased tau phosphorylation leads to the destabilization of the NMDAR:PSD95 and AMPAR:PSD95 complexes with increased AMPAR endocytosis resulting in reduced synaptic strength and LTP. Partial inhibition of MCI with CP2 increases AMP/ATP ratio that directly activates AMPK with concomitant effect on the activity of PP2Ac, CDK5 and GSK3β resulting in a significant reduction in pTau levels. Reduction in pTau and Fyn results in the stabilization of the NMDAR and AMPAR complexes and improved LTP leading to enhanced cognitive function in 3xTg-AD mice.

In this study, we specifically addressed whether treatment with MCI inhibitors attenuates pTau-related toxicity in AD. In our previous studies, we demonstrated that treatments with tool compound CP2 and other MCI inhibitors developed in our laboratory (data not shown) consistently reduced levels of soluble and insoluble Aβ and improved mechanisms associated with Aβ toxicity in APP, PS1 and APP/PS1 mice [48, 49]. The decrease in Aβ levels in these studies could be explained by the synergistic mechanisms where, as was shown previously, CP2 could disaggregate Aβ [78], while soluble peptides could be removed via the AMPK-dependent increase in autophagy [49]. Furthermore, similar to our findings in APP/PS1 mice [49], CP2 treatment significantly increased glucose uptake in the brain of male and female 3xTg-AD mice. Synaptic activity and LTP critically depend on ATP levels and a sustained availability of glucose for energy production in neuronal mitochondria [79]. Reduced glucose uptake and utilization in the brain detected using FDG-PET and peripheral glucose intolerance are present early in patients with mild cognitive impairment [10]. Recent reviews of drug discovery for neurodegenerative diseases of aging, including AD, identified strategies leading to energy restitution in the brain as the most promising to sustain cognitive function and combat neurodegeneration [79]. Both brain hypometabolism and peripheral glucose intolerance prominent in 3xTg-AD mice [56, 80] were reversed by CP2 treatment. In agreement with the published observations and similar to AD patients, vehicle-treated 16-month-old 3xTg-AD mice displayed a significant reduction in glucose uptake in the brain measured *in vivo* using translational biomarker FDG-PET. Compared to NTG mice, the decrease in glucose uptake in 3xTg-AD mice was significant, with the females being affected to a greater extent. Treatment with CP2 restored glucose uptake in male and female 3xTg-AD mice to the levels observed in NTG controls. Similarly, there was an improvement in peripheral glucose metabolism, specifically in females. Together with our previous studies showing that MCI inhibitors reduce Aβ pathology, the demonstration that this approach also alleviates pTau toxicity gives evidence of a promising role for MCI inhibitors in the prophylaxis and therapy of AD.

In our previous studies, CP2 treatment was conducted in female FAD mice only. Here we evaluated CP2 efficacy in male and female 3xTg-AD mice. While the most significant outcomes including a reduction of pTau levels, restoration of energy homeostasis in the brain, and cognitive protection were observed in male and female 3xTg-AD mice, females responded to CP2 treatment more consistently. Specifically, significant improvement of peripheral glucose metabolism was detected only in CP2-treated female 3xTg-AD mice. We also did not detect AMPK activation in CP2-treated male 3xTg-AD mice. At present, it remains unclear what parameters of the treatment paradigm might have contributed to these differences. It is feasible that the window of therapeutic opportunity, CP2 concentrations and/or the duration of treatment may need to be adjusted to achieve a better treatment response in males. However, it is also possible, based on the emerging evidence of sex-specific mechanisms of AD progression [81], that mitochondria-targeted therapeutics could be more beneficial in females in restoration of specific pathways that are differentially affected compared to males [82]. Further mechanistic studies should help to clarify these issues. The application of biomarkers that report on therapeutic efficacy and target engagement, including the recently identified blood/plasma levels of pTau181 [83], will also be considered in the future studies to address the observed sex-specific response to treatment.

Our mechanistic studies suggest that activation of AMPK plays a critical role in the neuroprotective mechanisms associated with mild MCI inhibition. While mechanisms involved in the AMPK-activated stress response are complex, inhibition of various components of mitochondrial electron transport chain, MCI in particular, using genetic or pharmacological manipulations, was shown to provide significant health benefits, increase longevity, protect against neurodegeneration, and improve mitochondrial function and cellular energetics in multiple model organisms [84-89]. Of key importance to the extension of health span in humans, correlation between mtDNA variability and longevity in a cohort of 2200 ultranonagenarians (and an equal number of controls) revealed that mutations in subunits of MCI that resulted in partial loss of its activity had a beneficial effect on longevity, while the simultaneous presence of mutations in complexes I and III and in complexes I and V appeared to be detrimental [90]. Additional support for the development of MCI inhibitors as disease-modifying strategy for neurodegenerative diseases and the safety of the application of MCI inhibitors in humans comes from metformin, an FDA approved drug to treat diabetes. Among other targets, metformin inhibits MCI. It is prescribed to the elderly population and has relatively safe profile even after chronic treatment [91]. Application of metformin has been shown beneficial in restoring energy homeostasis in the brain of APP/PS1 mice providing additional support for our findings [92]. Recent study conducted in a large Finish population of older people with diabetes demonstrated that long-term and high-dose metformin use do not increase incidences of AD and is associated with a lower risk of developing AD [93]. Resveratrol is another MCI modulator where its effect on MCI (activation or inhibition) depends on the concentration. It also activates PP2Ac reducing pTau levels in AD models [94]. Resveratrol is currently in clinical trials for multiple human conditions [95-97]. However, compared to CP2, these compounds have limitations associated with the lack of selectivity, specificity, and bioavailability justifying the need for further development of safe, specific and efficacious MCI inhibitors for clinical trials.

In conclusion, we provided the evidence that partial inhibition of MCI with a specific small molecule inhibitor had beneficial effect in 3xTg-AD mice. Chronic treatment was safe and resulted in reduced levels of pTau, improved brain energy homeostasis, LTP and synaptic activity leading to cognitive protection in male and female 3xTg-AD mice. Molecular mechanisms implicate activation of AMPK as an upstream singling pathway. Together with the previous studies, our data suggest that mitochondria represent a small molecule druggable target for AD treatment where restoration of synaptic function, LTP, energy homeostasis and cognitive protection could be achieved in the presence of Aβ and pTau –related pathologies.

## Acknowledgements

We thank Mayo Clinic Cores for help with the FDG-PET experiments. This research was supported by grants from the National Institutes of Health NIA RF1AG55549, NIA RO1AG062135, and ADDF 291204 (all to ET). Its contents are solely the responsibility of the authors and do not necessarily represent the official view of the NIH. The funders had no role in study design, data collection and analysis, decision to publish, or preparation of the manuscript.

